# Dicarbonyl stress enhances tumor intravasation

**DOI:** 10.1101/2024.07.03.601810

**Authors:** Nilesh Kumar, Bidita Samanta, KM Jyothsna, Varun Raghunathan, Prosenjit Sen, Ramray Bhat

## Abstract

Metastasis of cancer is a multi-step process that involves the migration of transformed cells from their native organ into a vascular channel, followed by their dissemination to prospective sites of colonization. The entry of tumor cells into blood or lymph, known as intravasation involves their breaching the stromal and endothelial extracellular matrix (ECM) and the endothelial barriers. How the kinetics of these cell-ECM interactions are confounded by chronic inflammatory stresses seen in comorbid risk factors of cancer such as diabetes and aging remain ill-investigated. Here, we construct and deploy a histopathology-motivated, imaging-tractable, microfluidic multi-organ-on-chip platform, that seamlessly integrates two tissue environments: that of a breast tumor and a vascular channel, to study the problem. The former comprises invasive triple-negative MDA-MB-231 breast cancer cells embedded within a three-dimensional fibrillar Collagen I milieu. The latter consists of a monolayer of TeloHAEC, immortalized human aortic endothelial cells arranged on laminin-rich basement membrane ECM, both of which concentrically line a hollow channel, wherein unidirectional fluid flows are implemented. The chip showcases the complexity of intravasation, wherein tumor cells and endothelia cooperate to form anastomotic structures. The formation of such structures is regulated by fluid flow in the vascular channel and is associated with cancer cell migration and entry into the vascular channel. Disseminated cancer cells are observed to enter, get adhered within, and flow through the vascular channel. Exposure to methylglyoxal (MG), a mediator of dicarbonyl stress associated with diabetic circulatory milieu, leads to greater cancer cell intravasation and flow through the vascular channel. This could be driven not just by MG-induced endothelial senescence and shedding, but also by the effect of MG on the chip ECM: we demonstrate it can degrade basement membrane and pathologically crosslink Collagen I, diminishing their cell adhesiveness. Our results thus show how dicarbonyl stress may attenuate homoeostatic barriers to cancer intravasation, exacerbating metastasis.

## Introduction

Metastasis of epithelial cancers typically involves the egress of transformed cells from their locus of origin into their surrounding connective tissue stroma, followed by migration to a local vascular fluid microenvironment [1]. Herein, single cells or disseminated tumor clusters migrate through blood or lymph and colonize favorable tissue- and organ-microenvironments, which allow the development of micro- and macro-metastatic foci. Each of these stages represent novel opportunities for the observation and analysis of intercellular (homotypic and heterotypic) and cell-extracellular matrix (ECM) interactions, which regulate the kinetics of invasion and metastasis [2–5].

The entry of cancer cells into the vasculature is referred to as intravasation [6]. Under active investigation, intravasation is thought to be guided by a variety of cues that include cell intrinsic alterations associated with tumorigenesis, such as the invocations of the transition from epithelial to mesenchymal states, migration-predisposing mutations, and metabolic reprogramming [7–9], changes in the ECM that facilitate a more directed migration of cancer cells toward vascular channels [10], and interplays between aberrant signaling and altered ECM mechanics that contribute to vessel directed invasion [11].

The integration into biological studies of advances in microscopic imaging and bioengineering has shed fresh light on tumor intravasation [12]. In vivo, an important method of gauging the ability of cancer cells to enter circulation is by measuring their numbers in blood [13]. Several techniques have been developed to image and measure the cancer cells entering the circulation through the utilization of their biophysical and biochemical properties [14–16]. Investigations using such techniques have introduced novel concepts such as the correlations of the presence in blood of clusters of disseminated cancer cells with greater metastatic efficiency [17]. However, given that the shear stresses within vascular compartments may destroy a significant proportion of disseminated cancer cells, a quantitative understanding of the efficiency of intravasation through the measurement of circulating tumor cells is difficult to arrive at. Furthermore, such measures do not provide a mechanistic understanding of the intravasation process. Such limitations can be overcome by a direct examination of intravasation using intravital two-photon microscopy within animal models [18]. However, perturbation studies probing biophysical interplays are often intractable through such experimental systems. Moreover, the molecular details of murine vascular anatomy can differ from that of its human orthologs necessitating complementary investigations using human cells.

Keeping such considerations in mind, on-chip platforms have reconceptualized investigations into such dynamic multistage biological processes such as tumor intra- and extra-vasation. Chips combining the cultivation of endothelia and cancer cells (with or without other stromal constituents) have allowed investigators to examine the contributions of intercellular interactions, secretome, and ECM remodeling to carcinomatosis [19–22]. Using this approach, a plethora of efforts have shed light on mechanisms of tumor intravasation, such as how cells traverse endothelial barriers through membrane extensions [23], how the matrisomal complexity regulates vascular entry of cancer cells [24], and the effect of flow on tumor dynamics [25]. Of metabolic alterations seen in cancer invasion, the role of dicarbonyl stresses is emerging as an important modulator of cancer progression [26]. Dicarbonyls are produced as byproduct metabolites of an excess glycolytic flux as seen in aged tissues or in diabetes mellitus [27]. By mediating glycation of proteins, lipids, and nucleic acids (and forming advanced glycation end products (AGE), they contribute to organ damage associated with diabetes such as nephro-, neuro-, retino-, and angio-pathy [28]. Although diabetes and age have been shown to be important risk factors for cancer incidence and metastasis, their specific cell biological associations with, or contributions to, the intravasation stage of cancer invasion remain ill-understood. Herein, we hypothesize that a debilitative effect on endothelial homeostasis by dicarbonyls could exacerbate tumor intravasation. We test this hypothesis using a customized tumor-vasculature chip that incorporates vascular-like flows using microfluidics and is tractable through wide field microscopic imaging and demonstrate that dicarbonyl stress can potentiate the vascular entry of invasive breast cancer epithelia.

## Results

### The chip recapitulates the histopathological milieu of the intravasation

In order to mimic a pathological context in which cancer cells inhabiting the breast cancer stroma enter a nearby blood vessel (Figure 1A), we designed our chip as a combination of a tumor microenvironment that was contiguous with a constructed vascular channel (Figure 1B) Figure 1C provides a schematic depiction of the macro- and microscopic design features and length scales of the chip components (see also the appropriate methods section for further details of the construction), and Figure 1D provides a 3D rendering of the continuity between the two chip compartments. Towards incorporating appropriate histopathological complexity, the vascular channel incorporated a monolayer of endothelia (depicted in green) overlaid on a thin layer of laminin-rich basement membrane (BM) (depicted in brown), with both these layers covering the entire interior of the channel leaving behind a central lumen for fluid flow (dotted black arrow) (Figure 1B). The tumor microenvironment chamber comprised breast cancer cells (depicted in red) embedded within a Collagen I stroma-like milieu (depicted as pink with fibers) (Figure 1B and E). Therefore, to intravasate, the cancer cells would need to traverse through Collagen I, breach the BM, move past the endothelia and then enter the lumen of the vascular channel.

**Figure 1:**
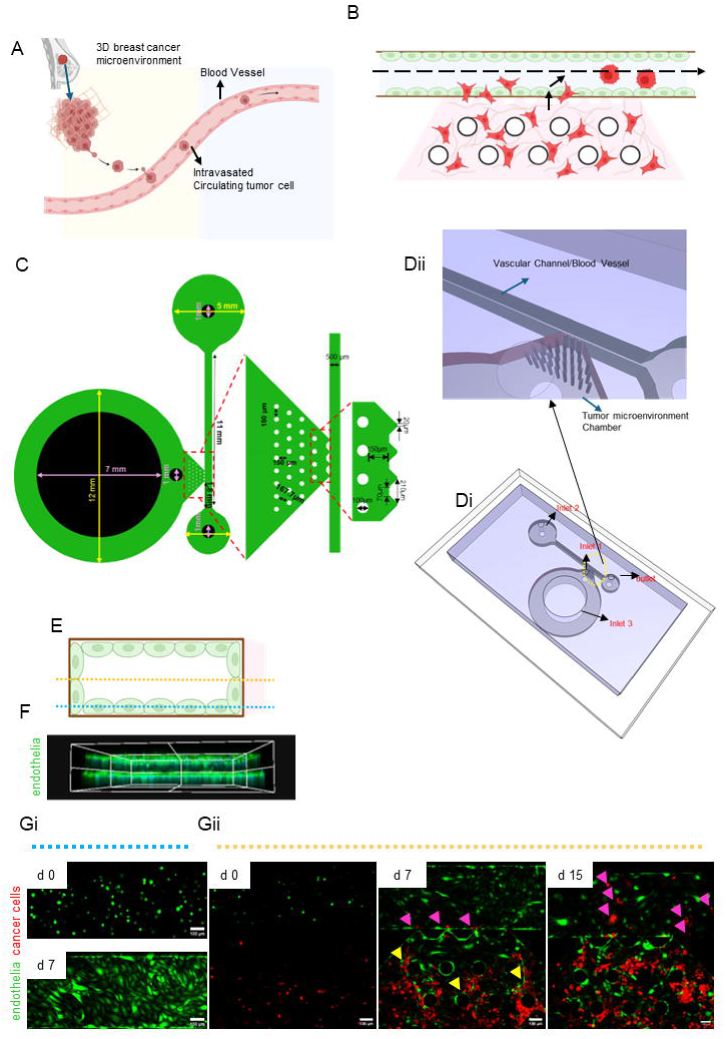
construction of an organ-on-chip (OOC) for studying intravasation of cancer cells.(A) is an anatomically inspired depiction of cancer cell intravasation. (B) schematically depicts the components of the intravasation-on chip platform comprising endothelia (green), endothelial casement membrane matrix (brown), Collagen I matrix (pink) and invasive breast cancer cells (red). The dotted arrow line indicates the flow of medium mimicking blood. (C) shows 2D schematic depiction of the device with detailed feature dimensions. Depicted leftwards is the inset of the interface of tumor microenvironment and vascular channel for circular pillars arrangement and trapezoidal pillars at the interface of two channels. (Di) shows the isometric projection of OOC device; inlet 1 is to introduce breast cancer cells embedded in Type 1 collagen. Inlet 2 and outlet are for the vascular channel inlet and outlet respectively. Inlet 3 is for the nutrient media reservoir to supply nutrients to 3D tumor microenvironment of the device. and (Dii) shows the isometric projection of device at the interface of tumor microenvironment chamber and vascular channel. (E) is a schematic depiction of the cross section of vascular channel shown in B. (F) Confocal photomicrograph showing the maximum intensity projection of TeloHAEC endothelial cells (expressing GFP; green) within the vascular channel. (Gi) Confocal micrographs showing the base of the vascular channel (digitally sectioned representative Z plane of the device shown with a dotted blue line in E) exhibiting attaching endothelial cells (green) on day 0 and a confluent monolayer of the cells by day 7. (Gii) Confocal micrographs showing the mid-section of the vascular channel and tumor chamber (digitally sectioned representative Z plane of the device shown with a dotted orange line in E) exhibiting endothelial cells (green) and triple negative breast cancer MDA-MB-231 cells (expressing RFP; red) on day 0 (left), day 7 (middle) and day 15 (right). Yellow arrowheads indicate areas where endothelial cells have entered the tumor chamber and formed anastomoses with the cancer cells. Pink arrowheads indicate breast cancer cells that have intravasated into the vascular channel. Scale bar for all figures = 100 µm.(A) and (B) schematics are prepared in BioRender

Under flow conditions maintained over 15 days, we confirmed the incorporation of TeloHAEC endothelia (stably expressing Green Fluorescent Protein) and MDA-MB-231 TNBC cells (stably expressing Red Fluorescent Protein) within their respective chambers within the chip on Day 0 (Figure 1Gi top and Figure 1Gii left; Gi shows the base of the vessel chamber at representative height of 1E shown by blue dotted line and Gii shows the mid-section of the vessel chamber of 1E shown by a yellow dotted line). By the 7^th^ day, we observed that the endothelia completely populated the vascular channel and had formed tightly apposed sheets (Figure 1Gi bottom). The cancer cells had also divided within the microenvironment chamber. Interestingly, the endothelial cells had penetrated within the microenvironment chamber and had organized themselves into anastomotic structures (yellow arrow heads) along with cancer cells with cellular rows comprising both cells forming a three-dimensional cavernous interface (Figure 1Gii middle). We also observed an infiltration of a small number of cancer cells into the vascular channel (purple arrow heads). By the 15^th^ day, we found a larger number of cancer cells had intravasated and had adhered to inner linings of the vascular channel (Figure 1Gii right, purple arrow heads). The channel-chamber interface was porous, suggesting bulk intravasation had physically breached the endothelial and matrix barriers.

### Vascular-cancer anastomoses are driven by flow and the presence of cancer cells

We next sought to uncover the factors that regulate the formation of the endothelial anastomoses within our device. To do so, chips were designed by incorporating only endothelia within the vascular channel and the unidirectional flow of medium was paused, although the latter was replenished regularly to prevent endothelial death due to lack of nutrients (Figure 2A left, a schematic depiction of the chip conditions. We observed endothelia were slow to populate the vascular channel and even on day 7, had not formed a confluent sheet at the base of the channel (base section view shown in Figure 2Ai middle, blue dotted line through the schematic depiction). Their penetration into the microenvironment chamber was considerably delayed compared with controls in Figure 1G (mid-section view shown in Figure 2Aii right, yellow dotted line through the schematic depiction).

**Figure 2:**
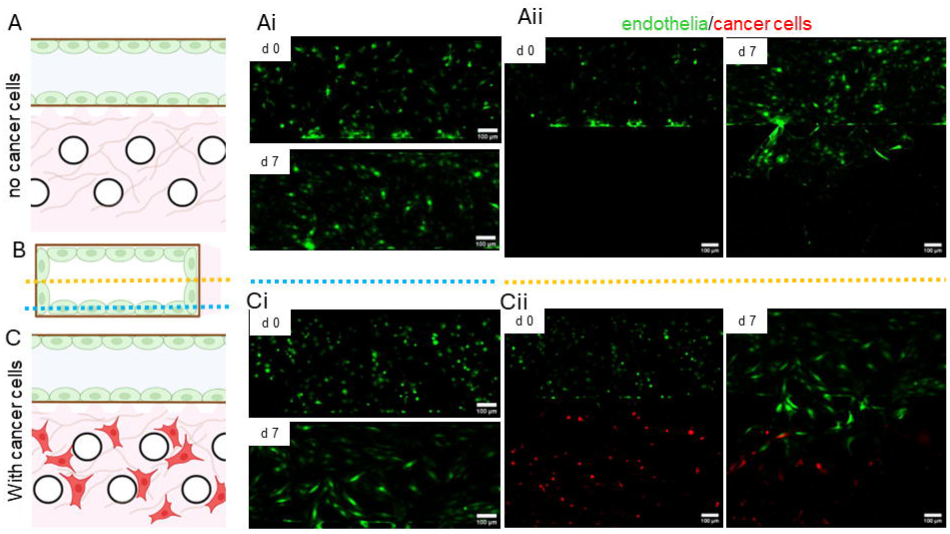
Intravasation rate is affected because of not enough supply of nutrient/media OR no flow of media through vascular channel condition. (A) schematically depicts the components of the intravasation-on chip platform comprising endothelia (green), endothelial casement membrane matrix (brown), Collagen I matrix (pink) and invasive breast cancer cells (red). The dotted arrow line indicates the flow of medium mimicking blood. (B) is a schematic depiction of the cross section of vascular channel shown in A. (C) Confocal photomicrograph showing the maximum intensity projection of TeloHAEC endothelial cells (expressing GFP; green) within the vascular channel. (Ai) Confocal micrographs showing the base of the vascular channel (digitally sectioned representative Z plane of the device shown with a dotted blue line in (B) exhibiting attaching endothelial cells (green) on day 0 and a confluent monolayer of the cells by day 7. (Aii) Confocal micrographs showing the mid section of the vascular channel and tumor chamber (digitally sectioned representative Z plane of the device shown with a dotted orange line in B) exhibiting endothelial cells (green) day 0 (left), day 7 (right). (Ci) Confocal micrographs showing the base of the vascular channel (digitally sectioned representative Z plane of the device shown with a dotted blue line in (B) exhibiting attaching endothelial cells (green) on day 0 and a confluent monolayer of the cells by day 7. (Cii) Confocal micrographs showing the midsection of the vascular channel and tumor chamber (digitally sectioned representative Z plane of the device shown with a dotted orange line in (B) exhibiting endothelial cells (green)) and triple negative breast cancer MDA-MB-231 cells (expressing RFP; red) day 0 (left), day 7 (right)

In the absence of flow when both endothelia and cancer cells were seeded into their respective chambers (schematic depiction shown in Figure 2C left), the spreading of endothelia in the lumen of the vascular chamber on day 7 was enhanced when compared with Figure 2A, even though flow was paused, although less than control (base section view shown in Figure 2Ci middle, blue dotted line through the schematic depiction). However, even on day 7 of culture we could not observe any cancer cells having intravasated or located close to the trapezoidal spatial continuities between the compartments. Although endothelia had penetrated the tumor chamber, the extent of their migration and interaction with cancer cells was lower than in control (mid-section view shown in Figure 2Cii right, yellow dotted line through the schematic depiction), suggesting that flow in the vascular channel is crucial for the migratory and interactive dynamics of endothelia allowing them to connect with cancer cells and also facilitating the latter to intravasate into the vascular channel.

### Exposure to methylglyoxal enhances intravasation

In order to mimic an inflammatory angiopathic vascular microenvironment, endothelial cells were treated with 200 μM methylglyoxal (MG), a glycolytic metabolite [29] that is known to be produced at high levels during metabolic stress [30–32], and which mediates the formation of advanced glycation end products within cells with known vasculopathic consequences [33–35] (setup is shown in Figure 3A). MG concentrations within cells have been measured to lie between 120 nm to 400 µM [36,37]. Kong and colleagues have estimated that treatment of cells with 100-500 µM MG would lead to an intracellular concentration of 100-700 nM, indicating our treatments are within physiological relevance [38]. We observed that the MG-treated cells attached to the base of the vascular channel but remained as patches by day 7 with large spread areas and a dysmorphic appearance, indicative of senescence (Figure 3C left compared with control Figure 3B left; yellow arrowheads).

**Figure 3:**
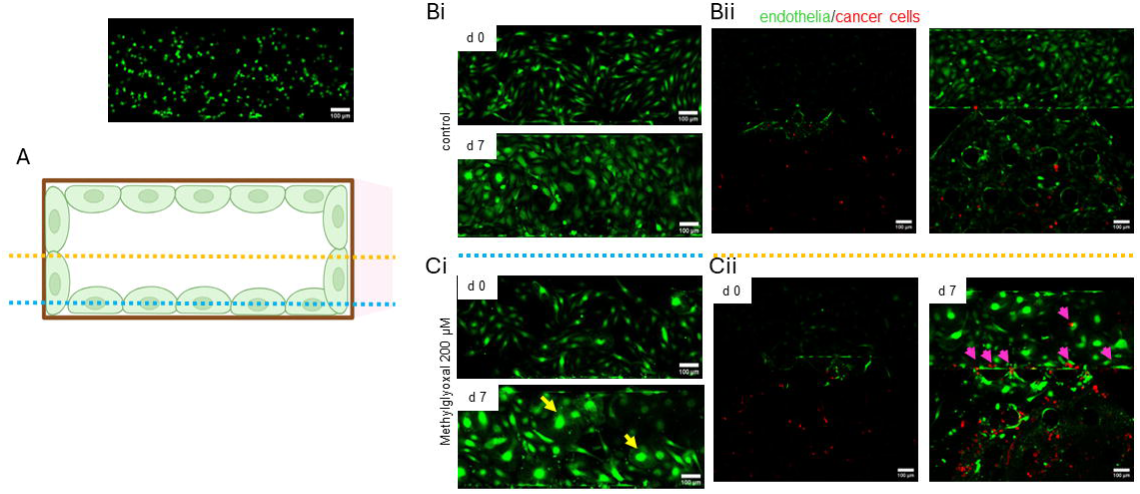
Intravasation in the case of endothelia affected by dicarbonyl stress. (A) is a schematic depiction of the cross section of vascular channel shown in (Fig2(A)). (Bi) Confocal micrographs showing the base of the vascular channel (digitally sectioned representative Z plane of the device shown with a dotted blue line in (A) exhibiting attaching endothelial cells (green) on day 0 and a confluent monolayer of the cells by day 7. (Bii) Confocal micrographs showing the mid section of the vascular channel and tumor chamber (digitally sectioned representative Z plane of the device shown with a dotted orange line in A) exhibiting endothelial cells (green) and triple negative breast cancer MDA-MB-231 cells (expressing RFP; red) on day 0 (left), day 7 (right). (Ci) Confocal micrographs showing the base of the vascular channel (digitally sectioned representative Z plane of the device shown with a dotted blue line in (A) exhibiting attaching endothelial cells (green) on day 0 and day 7. yellow arrowheads showing shredding of endothelia and senescent endothelia due to dicarbonyl stress. (Cii) Confocal micrographs showing the mid section of the vascular channel and tumor chamber (digitally sectioned representative Z plane of the device shown with a dotted orange line in B) exhibiting endothelial cells (green) and triple negative breast cancer MDA-MB-231 cells (expressing RFP; red) on day 0 (left), day 7 (right). Purple arrowheads showing intravasated cancer cells. Scale bar = 100 µm.

The penetration of endothelia treated with MG into the microenvironment chamber was slower than untreated controls (Figure 3Cii right compared with control Figure 3Bii right). Here too, the cells were found to be dysmorphic in appearance. To our surprise, even at day 7, several cancer cells were found to enter the vascular channel, where MG treatment was given, indicating the barriers were more porous and amenable to a more efficient intravasation of tumor cells (Figure 3Cii right compared with control Figure 3Bii right; purple arrowheads).

### Methylglyoxal increases crosslinking of Collagen I fibers

The kinetics of intravasation is known to be dependent on the interaction of cancer cells with both stromal cells and extracellular matrices [10,24]. Whereas the induction of senescence by MG of endothelia could debilitate their efficiency for barrier formation as has been observed through other modes of senescence induction in endothelia [39], thus allowing better cancer cell penetration into the vascular channel, we also asked if the metabolite had any effects on the structure and integrity of the ECMs used in the device. Therefore, different concentrations of MG (ranging from 50 µM to 200 µM) were added to 50 µL prepolymerized gels of Collagen I over 1 day and 4 days. The gels were subsequently stained using picrosirius red stain [40]. We observed a progressive decrease in intensity of the stain with an increase in the concentration and duration of MG (Figure 4A). The decreased intensity suggested either depletion of Collagen I due to its degradation by MG or a decreased availability of basic amino acid residues that could bind to the dye, which is consistent with the known crosslinking of lysines that is mediated by MG [41]. To investigate that the absence of staining was because of denaturation and dissolution of Collagen I, we examined the topography of collagen I scaffolds treated with MG using scanning electron microscopy (SEM). Photomicrographs showed the presence of Collagen I even after treatment with high concentrations of MG; however, a gradual attenuation in the fibrillar appearance was evident in scaffolds treated with progressively increasing MG concentrations compared with untreated controls (Figure 4B).

**Figure 4:**
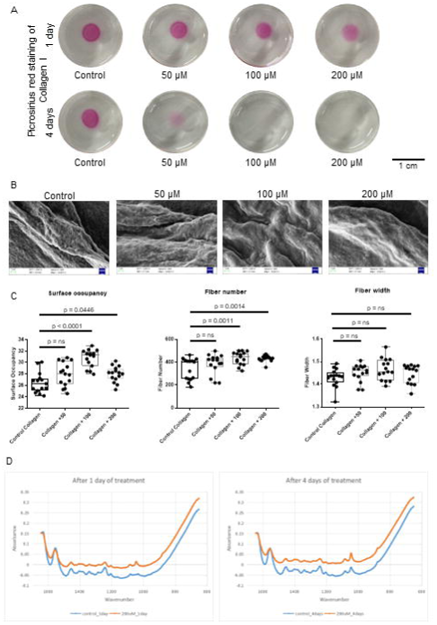
Methylglyoxal cross-links Collagen I (A) Photomicrographs of polymerized Collagen I gels, untreated (leftmost) and treated with increasing concentrations of 50 μM, 100 μM and 200 μM of methylglyoxal (left to right) for 1 day (top row) and 4 days (bottom row) and stained with picrosirius red (pink color). (B) Scanning electron micrographs of Collagen I gels untreated (leftmost) and treated with increasing concentrations of 50 μM, 100 μM and 200 μM of methylglyoxal (left to right) for 4 days (C) Scatter plot graphs showing surface occupancy (left), fiber number (middle) and fiber width (right) in Collagen I gels, untreated and treated with 50 μM, 100 μM and 200 μM of methylglyoxal for 1 day, imaged using second harmonic generation microscopy. (See also Supplementary figure 2) (D) Absorbance spectra of Collagen I gels untreated or treated with 200 μM methylglyoxal for 1 day (left) and 4 days (right) using Fourier Transform Infrared Spectroscopy (FTIR) under Attenuated Total Reflectance (ATR) mode. Scale bar for (A) = 1 cm. Scale bar for (B) = 1 μm.. One way ANOVA was performed for (C) on all concentrations and compared to control using Dunnett’s test.

That the Collagen I fibers were crosslinked was further indicated with the help of second harmonic generation (SHG) microscopy (Figure S2), wherein we observed an increase in fiber number, fiber width as well as surface occupancy of the fibers with a progressive increase in MG concentration (Figure 4C). To further confirm MG-induced Collagen I crosslinking, Fourier Transform Infrared Spectroscopy under the Attenuated Total Reflectance (FTIR-ATR) was performed for control collagen I samples treated with only PBS compared with those treated with 200 μM of MG. The absorbance versus wavenumber spectrum obtained for both the 1 day-samples and 4 days-samples were normalized on the amide I band, observed at 1645 cm^-1^ [42]. The spectra obtained was compared from 1300 cm^-1^ to 950 cm^-1^, called carbohydrate sugar band, consisting of the peaks at 1032 cm-1 and 1082 cm-1 [43]. The increased absorbance for 200 μM MG-treated sample compared to control sample, in this region can be attributed to increased presence of covalently linked MG within the Collagen I scaffolds (Figure 4D).

To investigate the effects of such fibrillar coalescence of Collagen I on cancer cells, adhesion was assayed on Collagen I gels, which were treated for 1 and 4 days with MG and then washed rigorously with PBS to prevent any confounding effect of residual MG on cells. Subsequently, both endothelial and cancer cells were added on the gels and their adhesion and spread morphologies studied. On MG-treated Collagen I, attached MDA-MB-231 and TeloHAEC cells were found to be circular rather than their mesenchymal spindle-shaped appearance (Figure 5Ai and Aii). In gels that were treated for 1 or 4 days with MG, there was a progressive decrease in adhesion of cells with an increase in the concentration of MG treatment (Figure 5Bi and Bii). This was clearly indicative of the decrease in density of adhered cells due to increased crosslinking that increases the difficulty of the cells to attach to the matrix.

**Figure 5:**
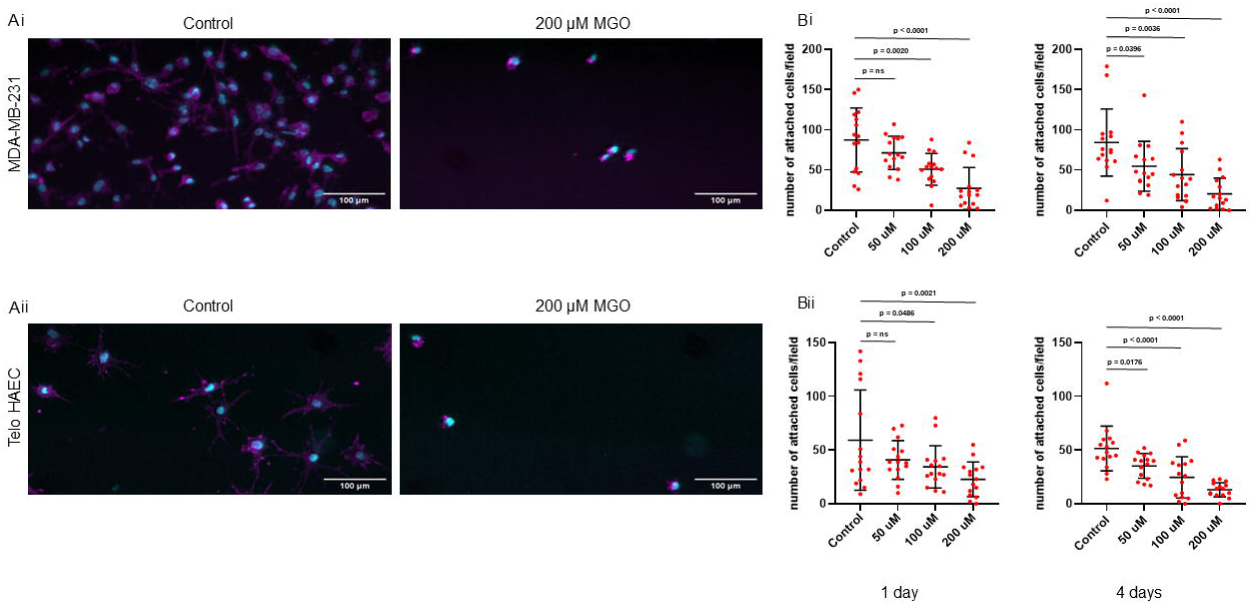
Methylglyoxal decreases cell adhesion on Collagen I (A) DAPI (cyan) and Phalloidin (magenta) stained images of cancerous cells MDA-MB-231 (top,i) and endothelial cells TeloHAEC (bottom,ii) adhered to collagen I gels, untreated (left) and 200 μM of Methylglyoxal (right) for 4 days (B) Scatter plot graphs for number of attached cells per field for 15 fields from 3 sets on Collagen I gels, untreated (leftmost) and treated with increasing concentrations of 50 μM, 100 μM and 200 μM of methylglyoxal (left to right) for 1 day (left) and 4 days (right). Scale bar for (A) = 100 μM. One way ANOVA was performed for (B) on all concentrations and compared to control using Dunnett’s test

### Methylglyoxal dissolves laminin-rich basement membrane and decreases cell adhesion

We next investigated the effects of MG on the LrBM matrix, which constitutes the ECM of the vascular channel. The chemical was added to different concentrations on 50 μL prepolymerized gels of LrBM for 1 and 4 days, following which they were stained for Alcian Blue to detect native mucopolysaccharides [44] (Figure 6A). We found not just that the staining was progressively lost with time of exposure and MG concentration, but that the gels too could not be visualized. To parse between whether this was a consequence of ECM disintegration or a leaching away of the glycan content, we stained the gels with crystal violet. We found an appropriate decrease in crystal violet staining with concentration and time of exposure (Figure 6B) in case of LrBM as compared to collagen I, indicating a denaturation and dissolution of the laminin with increased MG exposure.

**Figure 6:**
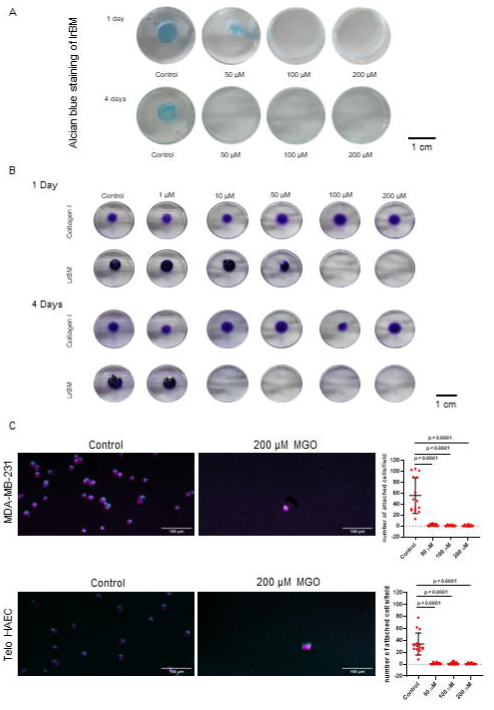
Methylglyoxal dissolves Laminin-rich Basement membrane and reduces cell adhesion on LrBM (A) Photomicrographs of polymerized LrBM gels, untreated (leftmost) and treated with increasing concentrations of 50 μM, 100 μM and 200 μM of methylglyoxal (left to right) for 1 day (top row) and 4 days (bottom row) and stained with Alcian blue (blue color) (B) Photomicrographs of polymerized collagen I and LrBM gels compared against each other using Crystal Violet staining, untreated (leftmost) and treated with increasing concentrations of 1 μM, 10 μM, 50 μM, 100 μM and 200 μM of methylglyoxal (left to right) for collagen I gels treated for 1 day, LrBM gels treated for 1 day, collagen I gels treated for 4 days and LrBM gels treated for 4 days (top to bottom) (See also Supplementary figure 3) (C) DAPI (cyan) and Phalloidin (magenta) stained fluorescent images for cancerous cells MDA-MB-231 (top) and endothelial cells TeloHAEC (bottom) adhered to LrBM gels, untreated (left) and treated with 200 μM of Methylglyoxal (middle) for 1 day and scatter plot graphs showing number of cells attached per field on LrBM gels for cancerous MDA-MB-231 (top-right) and endothelial cells TeloHAEC (bottom-right), untreated (leftmost) and treated with increasing concentrations of 50 μM, 100 μM and 200 μM of methylglyoxal (left to right) for 1 day. Scale bar for (A) = 1 cm, Scale bar for (B) = 1 cm and Scale bar for (C) = 100 μM. One way ANOVA was performed for scatter plot graphs of (C) on all concentrations and compared to control using Dunnett’s test.

After 1 day of treatment, Absorbance values of collagen I samples, untreated as well as treated with methylglyoxal, show consistency while absorbance values of LrBM samples show a significant decrease in case of 100 μM and 200 μM concentration as compared to untreated sample and samples with lower concentrations of methylglyoxal. For samples treated after 4 days, absorbance of collagen I still showed consistent values, whereas absorbance of LrBM showed significant decrease in values for 10 μM, 50 μM, 100 μM and 200 μM methylglyoxal treated samples as compared to untreated and 1 μM methylglyoxal treated samples. This indicates dissolution of LrBM gels with increased MG exposure as compared to increased crosslinking in case of collagen I gels. (Figure S3).

In cell adhesion studies carried out on LrBM treated with MG for 24 hours, endothelia and breast cancer cells were allowed to attach for 30 minutes followed by fixation and imaging using DAPI and Phalloidin: attached cells showed poorer spreading on MG-treated gels (Figure 6Ci and 6Cii). With an increasing concentration of MG treatment, both cells showed poorer ECM adhesion, consistent with observed poorer attachment of cells on plastic substrata than ECM.

## Discussion

Early events of cancer dissemination: migration of cells through stromal ECM, as well as late events such as colonization of prospective sites of metastasis by disseminated transformed cells have been extensively investigated using combinations of murine and organotypic human 3D culture models [45–47]. However, intermediate events such as intravasation remain poorly understood [48,49]. This is because the kinetics of intravasation is guided not just by the stromal microenvironment, where intravasation begins but also is dependent on the interactions between distinct cell populations, such as the connective tissues surrounding malignant tumors and vascular tissues which perfuse the former. It is to visually map the kinetics of intravasation that we constructed a histologically motivated on-chip device that allowed the physical interaction between a microenvironment chamber, which for our present study was loaded with breast stromal Collagen I and breast cancer cells, and a vascular channel, which had an outer layer of laminin-rich basement membrane matrix and an inner monolayer of endothelia.

It is within such histotypic environments that we were able to appreciate the flow- and cancer cell-driven migration of endothelia into the collagenous stroma, consistent with neoangiogenic morphogenesis seen during cancer invasion [50,51]. Using such a chip we were able to observe the effect of dicarbonyl stress specifically on the efficiency of tumor cell intravasation and found the latter to be higher in methylglyoxal-treated vascular environments compared with untreated controls. Methylglyoxal is a low abundance metabolite that accumulates within plasma of healthy individuals to concentrations of up to 300 nM and within cells up to 2 μM [52,53]. However, this may increase considerably in chronic inflammatory diseases that elevate glucose flux or impair the efficiency of detoxifying enzymes such as glyoxalases [54]. Therefore, the pathologic effects of MG are not limited to only diabetes, wherein accumulation of MG and AGE is being thought of as good if not better, a biomarker, but also in end stage kidney diseases and ageing or degeneration related organopathies [55,56]. We have used higher levels of MG in our experiments than would be seen in healthy individual serum, because we wished to mimic a longer cumulative exposure to low levels of dicarbonyls, which accumulate within tissues by covalently linking endogenous nucleophiles that are part of proteins and nucleic acids. It is impractical to subject experimental laboratory systems to exposure on time scales that are comparable to exposures within healthy individuals.

Although in our setup, endothelia are exposed for a day longer than breast cancer cells are to MG, it is interesting that the latter do not show any signs of senescence similar to the former. This could be attributable to the fact that MDA-MB 231 may have higher glyoxalase-GLO1 as indicated by high transcript levels in the Human Protein Atlas [57]. GLO1 and its allied enzymes convert MG into D-lactate by acting on the hemithioacetal formed by MG with glutathione (GSH) [58,59] D-lactate in turn can be converted to pyruvic acid by Lactate dehydrogenase D and participate in TCA cycle [60].

Under continuous flow, the senescent endothelia are shedded from the BM substratum. However, mechanobiological investigations suggest that senescence induction tends to increase matrix traction and adhesion through bigger focal adhesions and upregulated transcription of integrins [61,62]. We also observed the cancer cells to adopt a more amoeboid appearance in the MG-treated chips. This led us to ask whether the effects we were observing were a direct function of MG on the ECM rather than on the cells. Our observations, which show an exaggerated cross linking of Collagen I, suggest that the resultant depletion in amino acid moieties that could participate in binding cancer cells may lead to a mesenchymal-to-amoeboid transition in their migratory phenotype leading to an accelerated movement through stroma. ECM proteins tend to have high proportions of arginines and lysines which participate in ligand-receptor interactions for cell-matrix adhesion. Their reaction with MG and unavailability to mediate cell interactions may explain the phenotype we observe in our device. We also find that BM matrix on the other hand tends to get denatured, which could be explained by a weakening of the inter-ECM protein interactions by interfering covalent linkages by MG. The denaturation and dissolution of the endothelial ECM barrier may thus accelerate the shedding of endothelia and the intravasation of amoeboid cancer cells.

Despite the rich insights it affords, our chip is not without its limitations: the vascular channel is constructed in a manner that disallows rheological compression and relaxation. The vessel we construct approximates the anatomical complexity of capillaries, but it lacks pericytes. Moreover, its construction limitations constrain us to work with a design where the cross section of the vessel is rectangular rather than circular. Future iterations of the device will aim to overcome these problems and extend the scope of the device to other questions such as the effect of MG on extravasation, and cancer-cell endothelial adhesion.

## Materials and Methods

### Device Fabrication

The OOC device mold was fabricated using lithography and dry etching process at the nanofabrication facility at the Centre for Nanoscience and Engineering Department, Indian Institute of Science. Briefly, the OOC device design layout was created using the Clewin layout editor (WieWeb software). The photoresist (PR AZ-4562) was coated on a 4-inch Silicon wafer to fabricate a negative mold of the device on the Silicon wafer. The design was printed on PR using the Mask writer Heidelberg µPG501 tool. Subsequently, etching process and PR removal were performed using the DRIE Bosch process (SPTS DRIE). Subsequently, the device was fabricated using soft lithography with polydimethylsiloxane (PDMS) at a ratio of 10:1(w/w)::elastomer: crosslinker (Sylgard ^TM^ 184 Silicon Elastomer). The PDMS mixture was prepared and poured over the silicon wafer mold of the device and degassed to remove air bubbles in the mixture by placing it inside a desiccator using a vacuum pump. After degassing, PDMS was cured at 110^0^C for 20 minutes. Once cured, the PDMS slabs of the devices were cut, and holes were punched using 1mm diameter size biopsy punches for inlets 1,2, and outlet (Figure 1D). The 7mm diameter hole was punched for the media reservoir (inlet 3, Figure 1D) associated with the cancer chamber of the device. Parallelly, some PDMS blocks were cut, and 1mm and 4 mm dia. holes were punched using the biopsy punches for facilitating as a holder of microtubes and blocker for media reservoir during the flow, respectively. Processed PDMS slabs were bonded to cleaned glass slides (75X25 mm) using Harrick plasma cleaner at 100 watts, 0.4-0.6 bar, and an exposure time of 30-40 seconds. Prior to using the cells, devices were exposed to UV overnight for sterilization.

### OOC Device Details

The tumor chamber is circular with a diameter of 12 mm. It has a trapezoidal converging face towards the vascular channel. It is connected to the vascular chamber through 20 µm X 40 µm openings at four places by evenly spaced PDMS trapezoidal pillars. We introduced three notable features to the tumor chamber through iterative rounds for device engineering: (A) A 7mm opening (inlet 3, Figure 1(D)) at the top of the chamber, which can be filled with a nutrient-containing cell culture medium, allowing for the survival of cancer cells. This bigger opening enhances the static nutrient media storage capability to supply nutrients to the 3D tumor microenvironment. It allows additional benefits to users to change media frequently and addition of other biochemical /drugs without disturbing its structural integrity. During the iteration, we came up with the bigger static reservoir because a standard 1mm hole could not supply enough nutrients to the cancer cells, resulting in a hypoxia condition. The device does not need a separate neighboring channel to the 3D tumor microenvironment in the device to supply nutrients, showing the advantage of doing 3D cell culture in the device more conveniently. (B) In the design of the tumor chamber adjacent to the vascular channel, a diminutive 1mm inlet (inlet 1, Figure 1(D)) was engineered to facilitate the introduction of unpolymerized Collagen I, along with other hydrogels and oncogenic cells. This alteration permits the seamless integration of the cancer cell-laden Collagen I at the juncture between the tumor chamber and the vascular channel. Employing an additional 1 mm inlet hole (inlet 1) offers a distinctive advantage over the 7 mm hole (inlet 3) for introduction of cancer cells embedded Collagen I in the tumor chamber because, the smaller diameter ensures a more controlled and uniform injection of the cell-collagen matrix, overcomes the challenges posed by the inertia of the cells to get pushed to the interface of vascular channel. It also allows to preserve the structural integrity of 3D tumor microenvironment during seeding in the device and uniform distribution of cells. (C) In proximity to the vascular channel, an array of vertical PDMS pillars, each with a diameter of 100 µm, is organized into four rows. The rows are spaced 150 µm apart, and within each row, individual pillars are also separated by 150 µm (Figure 1(C), (D)). This configuration is designed for increased flow resistance against the flow in the vascular channel and to maintain the stability of the 3D tumor microenvironment, especially in the case of culture softer 3D tissue microenvironment. Additionally, the array of pillars offers significant adhesion properties for Collagen I, contributing to the formation of a stroma-like microenvironment. The zigzag pattern of the pillars across the columns introduces greater flow resistance against the shear experienced by the flow in the adjacent channel of the device, compared to a parallel arrangement. At the interface of tumor chamber and vascular chamber, a sequence of trapezoidal pillars, positioned with a minimum separation of 20 µm, supports both unicellular and collective cell migration processes.

The vascular chamber is designed as a rectangular parallelepiped channel, with dimensions of 500 µm in width, 40 µm in height, and 11 mm in length. It features an inlet with an expanded diameter of 5 mm and an outlet diameter of 3 mm (Figure 1(C), (D)). The outlet is positioned nearer to the tumor chamber and vascular channel interface than the inlet to minimize any perturbations to biologically significant interfacial phenomena that may arise from the turbulence of nutrient inflow. Considering the challenges associated with soft lithography processed with PDMS, the device depth is determined at 40 µm to ensure structural integrity and reproducibility.

### Cell Culture

For the tumor microenvironment, we used the triple negative breast cancer cell line MDA-MB-231 (ATCC) that was made to stably express red fluorescent (RF) using lentiviral transduction. The MDA-MB-231-RFP cells were cultured in DMEM:F12 (1:1) medium (HiMedia, AT007F) supplemented with 10% fetal bovine serum (Gibco, 10270). MDA-MB-231 cells were trypsinized using a 1:5 dilution of 0.25% trypsin and 0.02% EDTA (HiMedia, TCL007). After trypsenizing we used 1×10^6^ cells diluted in 60 µl of DMEM-defined medium (composition: DMEM:F12, Insulin, Hydrocortisone, Sodium Selenite, Estradiol, transferrin). Simultaneously, we neutralized acid-extracted Collagen I (3 mg/ml, Gibco, A1048301) using 10X DMEM and 2N NaOH; It was diluted to a concentration of 1 mg/ml with DMEM-defined medium containing MDA-MB-231 cells.

We prepared TeloHAEC endothelial cells (ATCC) stably expressing green fluorescent (GF) using lentivirus transduction for the vascular channel. TeloHAEC (endothelia) was cultured in EBM-2 Media (Lonza, #CC3302). Following the same protocol as for the MDA-MB-231 cells, we trypsinized the endothelial cells and used a cell density of 1×10^6^ cells in 40 µl for the vascular channel in the device.

### Cell seeding in OOC device

In the OOC device, we incorporated MDA-MB-231 cells embedded in Collagen I (1 mg/ml) for 3D tumor microenvironment. Following the preparation of these cells embedded in Collagen I, we introduced them into the device through inlet 1 (as shown in Figure 1(D)). Subsequently, the device was incubated at 37^0^C and 5%CO_2_ for 30-45 minutes to allow for Collagen I polymerization. Once the Collagen I polymerized, we introduced DMEM-defined medium in inlet 3 (Figure 1(D)) and into the adjacent straight channel by inlet 2 designed for the vascular channel. To facilitate the slow perfusion of nutrients to the cancer cells from this adjacent channel, we placed drops of DMEM-defined media (200 µl and 100 µl) at the inlet and outlet of the straight channel respectively.

The device was kept in the incubator at 37^0^C and 5%CO_2_ for 2-3 hours for MDA-MB-231 cell attachment on Collagen I fibers. Next, we introduced a laminin-rich basement membrane (LrBM; Matrigel^TM,^ Corning, 354230) into the straight channel of the device designed for the vascular channel through inlet 2. LrBM was diluted to a concentration of 100 µg/ml in DMEM-defined medium to provide a basement membrane for the endothelial cells to mimic the biological condition of blood capillaries. The device was then kept in the incubator at 37^0^C and 5%CO_2_ for an additional 4 hours to ensure complete coating of the LrBM on the vascular channel’s wall.

TeloHAEC cells were trypsinized and diluted to the specified concentration. After trypsenizing, endothelial cells were introduced into the device by inlet 2, and images were taken using the Olympus FLUOVIEW^TM^ FV4000 Confocal Laser Scanning Microscope for the subsequent time points.

### Perfusion System

To investigate endothelial attachment on the basement membrane, we maintained the device under static conditions for 24 hours. During this period, a static reservoir droplet of 100 µl was placed at the inlet and outlet of the device, and the reservoir was changed every 4 hours.

Next, we established a flow setup using the static head of the nutrient media reservoir. This static head was created using two nutrient media glass bottles (Figure S-1). One bottle was filled with 40 ml of EBM-2^TM^ media, serving as the supply reservoir, while the other bottle remained empty, acting as the sink. We connected microtubing (BD Intramedic^TM^, 427406) with an inner diameter of 380 µm from the supply bottle to the inlet of the vascular channel within the device. The height of the supply reservoir was adjusted to achieve the desired flow rate (approximately 0.5-1 ml/hr).

To monitor the flow rate, we observed the collection at 6-hour intervals. The initial observation images were taken at T = 4 days (d 4), followed by subsequent observations at T = 7 days (d 7), 10-11 days, and 14-15 days (d 15).

### Experiment method details for MG treatment

In a 60 mm Petri dish, 1.6×10^4^ cells were initially seeded. After 24 hours, 200 µM MG (M0252, Sigma-Aldrich) was added to the cell culture media (EBM-2 ^TM^) to pre-expose the cells to dicarbonyl stress. Once the cells reached 70% confluency, they were passaged and diluted with 40 µl of cell culture media before being seeded in the experimental device. The flow setup was established at T = 24 hours using the cell culture media containing 200 µM MG.

### Study of Methylglyoxal treatment on Collagen I

Collagen I gels preparation The stock of Collagen I, with the concentration of 3 mg/ml, was neutralized with the help of 2N Sodium Hydroxide (NaOH), 10X DMEM, Dulbecco’s Modified Eagle Medium (HiMedia, AT006F) and 1X PBS (Phosphate buffered saline) to a working concentration of 1 mg/ml. The neutralized Collagen I was utilized in preparing gels of 50 μL each for staining and microscopy experiments and coats of 100 μL each for cell-based experiments. The gels were allowed to polymerize at 37^0^C for 30 minutes. The polymerised gels were further treated.

### Picrosirius staining

Picric acid solution of 10 ml was prepared by adding 0.12 g of Picric acid into a beaker containing 10 ml of distilled water. The contents of the beaker were mixed well and then heated for about 5-10 minutes on a hotplate. This was followed by addition of 0.01 g of Direct Red 80 (Sigma Aldrich, 365548-5G) on the solution and mixed well to get 10 ml of Picric acid solution. The Collagen I gels, that were incubated in 1X PBS (control, untreated), 50 μM, 100 μM and 200 μM methylglyoxal for 1 day and 4 days, were washed thrice with 1X PBS and then with distilled water after the completion of their incubation. Picric acid solution was added on top of the gels and was stored for 2 hours. This was followed by washing the gels in 0.05% of acetic acid and absolute alcohol. The gels were washed thrice again with 1X PBS and stored in 1X PBS for 24 hours to get rid of the excess stain. The gels were then imaged.

### Scanning Electron Microscopy

Collagen I gels of 50 μL were washed thrice in 1X PBS after incubation in 1X PBS (Control), 50 μM, 100 μM and 200 μM methylglyoxal for 1 day and 4 days. The gels were then fixed in 1% formaldehyde (Thermo Fisher Scientific, 24,005) for 12-24 hours. After fixation, they were washed in distilled water for 5 minutes, followed by washes of 50% ethanol, 75% ethanol, 90% ethanol and thrice in 100% ethanol for 15 minutes. The gels were then air dried at room temperature and gold sputtered with the help of Jeol Smart Coater with the samples at 20 mm working distance for 1.5 minutes, resulting in a coat of approximately 5 nm. The gels were then imaged in ULTRA55 ultra high resolution scanning electron microscope integrated with energy and angle selective backscattered electron (EsB) detector. The images were taken at a working distance of approximately 9.7 mm and EHT of 5 kV at a magnification of 10 kX with the help of SE2 detector signal.

### Second Harmonic Generation Microscopy

The 50 μL collagen I gels were incubated in 1X PBS (untreated, control) and 50 μM, 100μM and 200 μM of methylglyoxal for 1 day. Methylglyoxal was removed after 1 day and the samples were washed thrice with 1X PBS to remove residual MG. The samples were then imaged using Second Harmonic Generation microscopy that involved a nonlinear optical imaging setup, utilizing a femtosecond fiber laser (Fidelity HP) emitting at 1040 nm with a pulse width of 140 fs and a repetition rate of 80MHz as the excitation source. The incident beam at fundamental wavelength was focused onto the collagen gels, which was mounted on an Olympus IX73 inverted microscope equipped with a ×60/1.2NA water immersion objective. The SHG signal at 520 nm emitted from the sample was collected using the same objective in an epi-detection setup. A dichroic mirror separated the fundamental and the second harmonic signal, and the SHG signal was detected using a photomultiplier tube (Hamamatsu, R3896) with band-pass (520 ±20 nm) and short-pass (890 nm cut-off) filters. SHG images were acquired by scanning the laser beam with a pair of galvanometric mirrors (Thorlabs, GVS002). A motorized sample stage (Thorlab, MLS203-1) was used to image a minimum of 15 different locations or fields of view (FOVs) sized at 50 x 50 μm² from each sample.

### Fourier transform infrared spectroscopy (FTIR) under Attenuated Total Reflectance (ATR) mode

The 50 μL Collagen I gels were incubated in 1X PBS (control) and 200 μM of methylglyoxal for 1 day and 4 days. Methylglyoxal was removed after 1 day and 4 days respectively and the samples were washed thrice with 1X PBS. 1X PBS was removed after washing and the gels were dried up. The infrared spectrum of Absorbance versus Wavenumber of the gels were obtained using FTIR spectrometer in UATR mode, containing detectors TGS and Liquid nitrogen cooled MCT, at a spectral resolution of 4 cm-1 for 32 scans through a Diamond crystal. The FTIR - Universal ATR spectrometer was atmosphere corrected, baseline and offset corrected and vector normalized. The graphs obtained were normalized for Amide I peak obtained at 1645 cm-1 and compared from 1300 cm^-1^ to 650 cm^-1^.

### Cell adhesion study on Methylglyoxal treated Collagen I

The 100 μL coats of Collagen I prepared on an eight-well chamber were incubated in the presence of PBS for control, 50 μM of MG, 100 μM of MG, and 200 μM of MG for 1 day and 4 days. MG was replenished after two days for the four-days samples. After incubation, MG was removed, and the coats were washed with 1X PBS thrice. Breast cancer cells MDA-MB-231 and TeloHAEC, that were cultured previously, were added to the coats such that each well contained approximately 10,000 cells. Cells were allowed to attach to the control and the MG-treated coats for 3 hours in an incubator at 37^0^C at 5% CO_2_ conditions. Cell suspension was removed from the wells after 3 hours and the cells that had attached were fixed using 4% formaldehyde for 30 minutes at 4^0^C. Cells are stored in 1X PBS at 4^0^C till further processing.

The fixed cells were stained for DNA using DAPI (D1306; Thermo Fisher Scientific) and F-actin using Alexa Fluor 568 Phalloidin (Invitrogen, A12380) for TeloHAEC and Alexa Fluor 488 Phalloidin (Invitrogen, A12379) for MDA-MB-231. The cells were permeabilized using 0.5% Triton X-100 (MB031, HiMedia) for 2 hours at room temperature, followed by wash with 0.1% Triton X-100 in 1X PBS thrice. MDA-MB-231 cells were incubated overnight with Alexa Fluor 488 Phalloidin dissolved in 0.1% Triton X-100 (1:500) at 4^0^C in the dark while TeloHAEC cells were incubated overnight with Alexa Fluor 568 Phalloidin dissolved in 0.1% Triton X-100 (1:500) at 4^0^C in the dark. The cells were then washed again thrice with 0.1% Triton X-100 for 5 minutes each. Both the cell lines were then incubated with DAPI dissolved in 0.1% Triton X-100 at 1:1000 dilution for 5 minutes in dark at room temperature. They were then washed thrice in 1X PBS. Images of the adhered cells were captured at 10X magnification in an Olympus IX83 inverted fluorescence microscope fitted with Aurox structured illumination spinning disk setup. The images were processed and cell counting analysis was performed for three biological replicates with 5 fields of view for each replicate using ImageJ software [63].

### Study of Methylglyoxal treatment on Laminin-rich Basement membrane (LrBM)

#### Laminin-rich Basement membrane gels preparation

Matrigel, consisting of Collagen IV, Laminin, Entactin and Percelan, of the stock concentration 7.6 mg/ml was directly added in the form of spherical gels of 50 μl for staining experiments and coats of 100 μL for cell-based experiments. The gels were allowed to polymerise at 37^0^C for 30 minutes before they were treated.

#### Alcian Blue staining

1X PBS (control), 50 μM MG, 100 μM MG and 200 μM MG were added to the collagen I gels of 50 μL and incubated for 1 day and 4 days. They were given a 1X PBS wash after incubation and fixed using 1% formaldehyde for 30 minutes at 37^0^C. They were again washed in 1X PBS and treated with 3% acetic acid solution for 3 minutes. This was followed by treatment with Alcian Blue stain, that was prepared by adding 1% w/V Alcian blue 8GX (HiMedia, TC359-10G) in 3% acetic acid solution at pH 2.5, for 30 minutes at 37^0^C. The gels were washed thrice with 1X PBS. Images were taken immediately after the final washing step.

#### Crystal violet staining

The 50 μL collagen I gels and LrBM gels were incubated with 1X PBS (control), 50 μM MG, 100 μM MG and 200 μM MG for 1 day and 4 days. After incubation was completed, MG was removed and the gels were washed twice with 1X PBS. LrBM gels were fixed using 1% formaldehyde for 30 minutes at 37^0^C and then again washed with 1X PBS to remove residual formaldehyde. Crystal violet stain of stock concentration was then added to the collagen I gels and LrBM gels for 15 minutes. Stain was removed and the gels were washed thrice with 1X PBS. Images were taken immediately after 1X PBS wash. After imaging, 1X PBS was removed from the samples and the crystal violet stained samples were stored for 24 hours for drying. The dried up samples were treated with methanol and kept at rocker for an hour to dissolve the crystal violet stain. 50 μL of the dissolved samples were added onto an assay plate with 150 μL of distilled water and mixed properly to dilute it to 1:4 ratio. The absorbance of the samples were measured at 595 nm using Tecan Infinite MPlex plate reader.

### Cell adhesion study on Methylglyoxal treated Laminin-rich Basement membrane (LrBM)

The 100 μL LrBM coats were prepared on an eight-well chamber. The coats were incubated for 1 day in 1X PBS (control) and 50 μM, 100 μM and 200 μM of methylglyoxal. After 24 hours, the treatment was removed and the coats were washed in 1X PBS thrice to get rid of the unreacted MG. MDA-MB-231 and TeloHAEC cells cultured were added onto the LrBM gels such that each well contained 10,000 cells. The cells were allowed to adhere to the LrBM matrix for 30 minutes in an incubator at 37^0^C at 5% CO_2_ conditions. The cell suspension was removed after 30 minutes from the wells and cells adhered to the coats are fixed using 4% formaldehyde for 30 minutes at 4^0^C. Cells are stored in 1X PBS at 4^0^C till further processing.

The fixed cells were stained for DNA using DAPI (D1306; Thermo Fisher Scientific) and F-actin using Alexa Fluor 568 Phalloidin (Invitrogen; A12380) for TeloHAEC and Alexa Fluor 488 Phalloidin (Invitrogen; A12379) for MDA-MB-231. The cells were permeabilized using 0.5% Triton X-100 (MB031; HiMedia) for 2 hours at room temperature, followed by wash with 0.1% Triton X-100 in 1X PBS thrice. MDA-MB-231 cells were incubated overnight with Alexa Fluor 488 Phalloidin dissolved in 0.1% Triton X-100 (1:500) at 4^0^C in the dark while TeloHAEC cells were incubated overnight with Alexa Fluor 568 Phalloidin dissolved in 0.1% Triton X-100 (1:500) at 4^0^C in the dark. The cells were then washed again thrice with 0.1% Triton X-100 for 5 minutes each. Both the cell lines were then incubated with DAPI dissolved in 0.1% Triton X-100 at 1:1000 dilution for 5 minutes in dark at room temperature. They were then washed thrice in 1X PBS. Images of the adhered cells were captured at 10X magnification in an Olympus IX83 inverted fluorescence microscope fitted with Aurox structured illumination spinning disk setup. The images were processed and cell counting analysis was performed for three biological replicates with 5 fields of view for each replicate using ImageJ software.

## Acknowledgements

This work was supported by the Wellcome Trust/DBT India Alliance Fellowship/Grant [IA/I/17/2/503312] awarded to RB. It was also supported by the John Templeton Foundation (#62220) and the Indo-French Centre for the Promotion of Advanced Research (69T08-2) to RB. NK and JKM acknowledge support from the Ministry of Education, Government of India. BS acknowledges support from the Prime Ministers Research Fellowship. We thank MeITY, Government of India for enabling the usage of micro and nano fabrication facilities in CeNSE, IISc. The opinions expressed in this paper are those of the authors and not those of the John Templeton Foundation.

Figure S-1 : (A) Macroscopic view of the device (B) Medium upon flow through our device was collected and collected material was cultured. (C) Medium collected at 7 days from control setups showed no cultured cells (2 representative epifluorescent micrographs) However, in 14 day collected medium, the two representative epifluorescent micrographs show attached cancer cells (yellow arrow heads) and debris of anoikic endothelia (red arrowhead). Scale bars = 100 µm

Figure S-2: Photomicrographs representing the fibers in polymerized collagen I gels, untreated (leftmost) and treated with 50 μM, 100 μM and 200 μM of MG (left to right) for 1 day, imaged using second harmonic generation microscopy.

Figure S-3: Bar graphs for polymerized collagen I and LrBM gels compared against each other using Crystal Violet staining, untreated (leftmost) and treated with increasing concentrations of 1 μM, 10 μM, 50 μM, 100 μM and 200 μM of methylglyoxal (left to right) for (i) collagen I gels treated for 1 day and LrBM gels treated for 1 day (ii) collagen I gels treated for 4 days and LrBM gels treated for 4 days.

