## Supplementary figures and images for "Dicarbonyl stress enhances tumor intravasation"

### Figure S-1

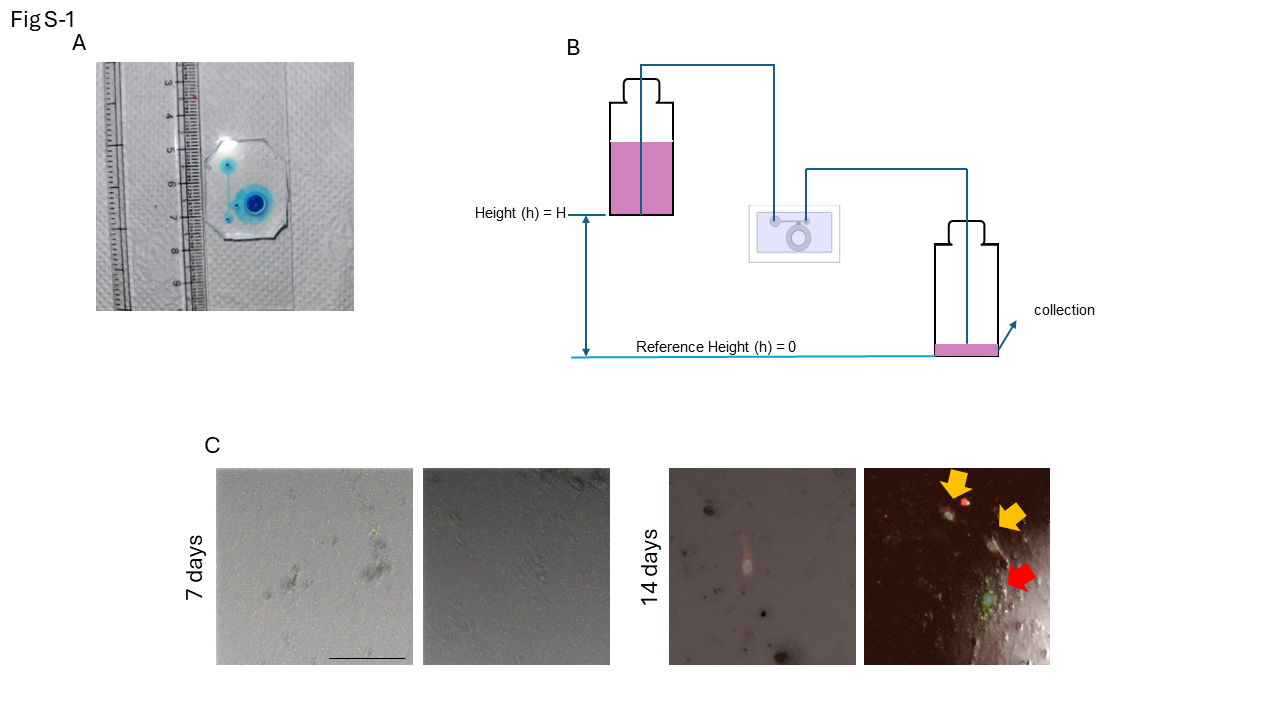

### Figure S-2

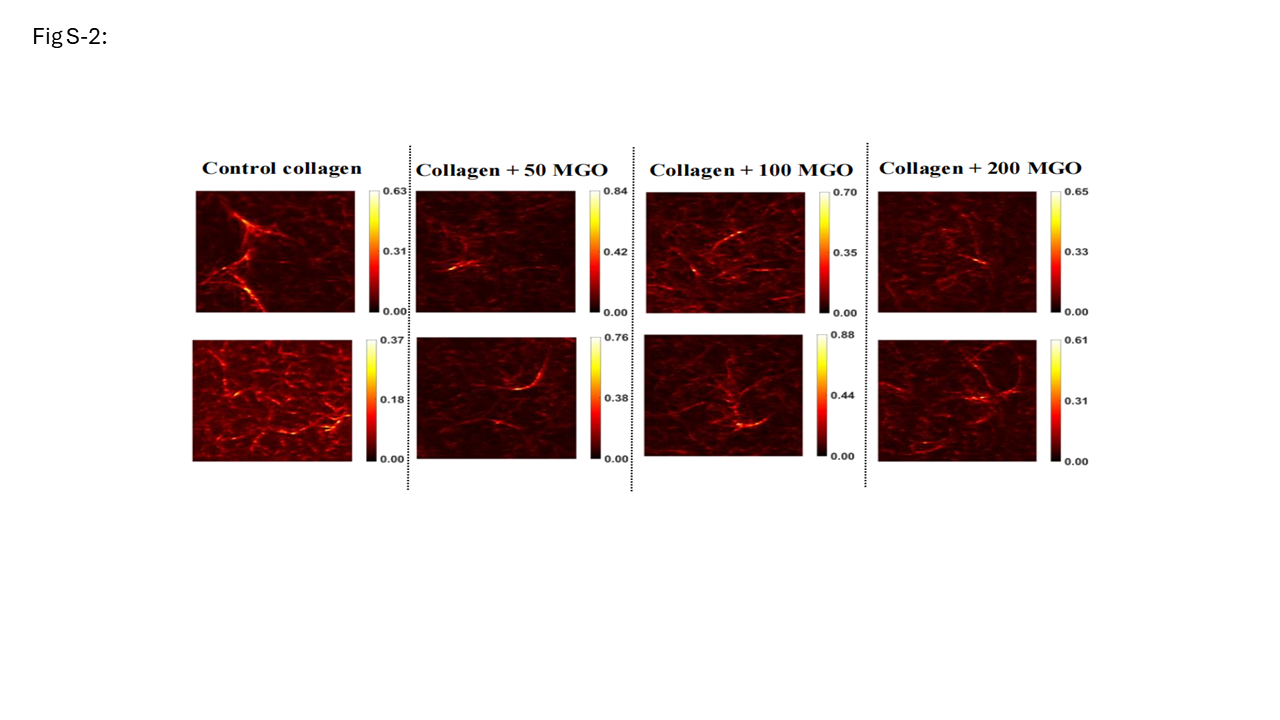

### Figure S-3

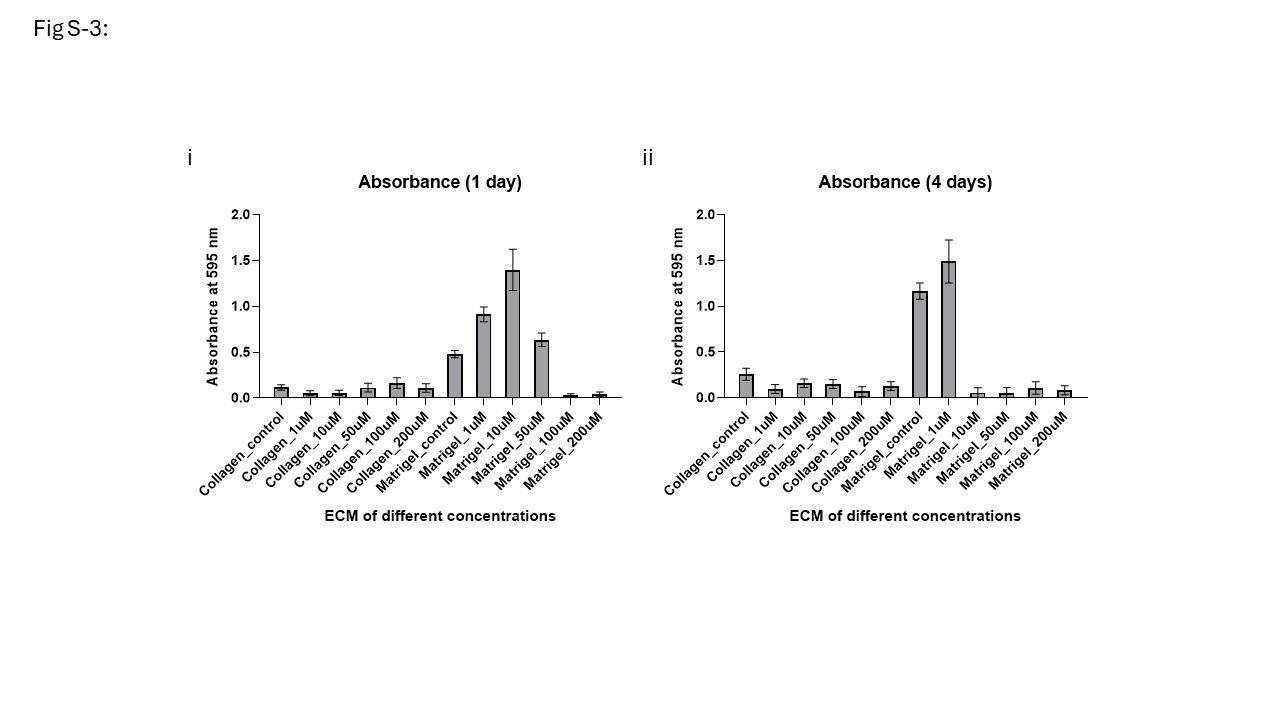
